# Activation of prefrontal cortex and striatal regions in rats after shifting between rules in a T-maze

**DOI:** 10.1101/2022.12.07.519425

**Authors:** Hongying Gao, Virginie Oberto, Ana Biondi, Susan J. Sara, Sidney I. Wiener

## Abstract

Prefrontal cortical and striatal areas have been identified by inactivation or lesion studies to be required for behavioral flexibility, including selecting and processing of different types of information. In order to identify these networks activated selectively during acquisition of new reward contingency rules, rats were trained to discriminate orientations of bars presented in pseudo-random sequence on two video monitors positioned behind the goal sites on a T maze with return arms. A second group already trained in the visual discrimination task learned to alternate left and right goal arm visits in the same maze while ignoring the visual cues still being presented. In each experimental group, once the rats reached criterion performance, the brains were prepared after a 90 minute delay, for later processing for c-fos immunohistochemistry. While both groups extinguished a prior strategy and acquired a new rule, they differed by the identity of the strategies, and previous learning experience. Among the 28 forebrain areas examined, there were significant increases in the relative density of c-fos immunoreactive cell bodies after learning the second rule in prefrontal cortex cingulate, prelimbic and infralimbic areas, in dorsomedial striatum and the core of nucleus accumbens, in ventral subiculum, and the central nucleus of the amygdala. These largely correspond to structures anatomically identified by previous inactivation studies. The data suggest that this dynamic network may underlie reward-based selection for action, a type of cognitive flexibility.

## Introduction

For survival, animals must adapt their behavior to changes in the external environment and to internal states. In goal-directed behavior, this can involve ignoring established stimulus reinforcement contingencies and shifting attention to a previously inconsequential stimulus. Previous work has shown that lesions of medial frontal cortex lead to impairments in shifting between rules in rats (Joel *et al*., 1997; Ragozzino *et al*., 1999a,b). Birrell and Brown (2000) demonstrated that lesions of the prelimbic (PL) and infralimbic (IL) regions of medial frontal cortex lead to impairment in shifting between strategies informed by different sensory modalities, (‘extra dimensional shifts) with no impairment in acquisition or reversal learning.

Elucidating the neural mechanisms underlying learning to select among alternative rules may involve different approaches including simultaneous recording from multiple brain regions in animals performing cognitively demanding tasks (e.g., Bissonette and Roesch, 2013; Oberto, et al., 2021). However, recordings during behavior can only provide correlative evidence. Task-related neural activity can only be considered necessary for execution of the task when inactivation of the structure recorded is shown to impair performance. Nonetheless, interpretation of inactivation studies is also limited since the vicarious action of multiple overlapping and parallel networks in the brain can permit normal performance even when a vital structure is inactivated. Complementary to the latter approaches are techniques measuring the activation of brain areas. C-fos, an immediate early gene product that is a metabolic marker for neural activity (Dragunow and Faull, 1989), can be a powerful tool to identify brain regions and networks that are differentially activated for distinct behavioral contingencies (see, e.g., Tronel & Sara, 2002). Here we identify multiple brain regions activated by an attentional shift, using immunohistochemical marking of c-fos.

On the basis of previous studies, our analyses focused on prefrontal cortical areas, their striatal projection zones, limbic areas including amygdala and dorsal and ventral hippocampal subfields as well as brainstem neuromodulatory centers (Birrell and Brown, 2000; Burnham, et al., 2010). A particular motivation for the present work is the observation that once rats learned and thus reliably received rewards for performing a behavioral strategy in a binary choice task, prefrontal cortical pyramidal neurons shifted their firing phase relative to hippocampal theta oscillatory activities (Benchenane et al., 2010). Furthermore, after learning, prefrontal interneurons more effectively inhibited principal neurons, and synchronously active prefrontal neuronal assemblies appeared for the first time (Peyrache, et al., 2009). These network changes are associated with learning of changes in response-reward contingencies. When this involves shifting attention to different cue types or sensory modalities, it can be referred to as an extra-dimensional shift. Lesions and pharmacological manipulation studies indicate that PL is vital for such learning (Birrell & Brown, 2000; Bissonette & Roesch, 2017). The present study examines c-fos immunoreactivity to identify the extent of the network reorganization related to learning of a new behavior-reward contingency.

## Materials and methods

Ten male Long-Evans rats (250-300 g, Janvier, Le Genest-Saint-Isle, France) were housed singly in cages under a 12 hour light-dark cycle. All rats were weighed and handled daily after their arrival. They had free access to water at first, and then were restricted to 10 min per day to motivate them to search for liquid rewards during pre-training and training days.

The T-maze (Figure 1A) was constructed from wood and painted matte black. The central stem and top alley were 1 m long and 8 cm wide with 2 cm high borders. The maze was elevated 70 cm above the floor and surrounded by a black cylindrical curtain 3 m in diameter running from floor to ceiling. Visual cues were displayed on two TV screens (80 cm diagonal) located behind the reward arms. When the rat crossed a photodetector beam on the central arm, vertical bars were displayed on one and horizontal bars on the other in a pseudo-random order (spatial frequency of 0.13 cycles per degree, viewed from the trigger point). Each of the 16 possible left-right sequences of visual cues was displayed at equal incidences over consecutive series’ of four trials, with the exception of three repetitions of the same cue to the left or the right (thus 14 sequences were used). Following correct choices, saccharinated (0.25%, 30 μl) water rewards were dispensed from small wells operated by solenoid valves controlled by a CED Power 1401 system (Cambridge Electronic Design, Cambridge, UK).

**Figure 1.**
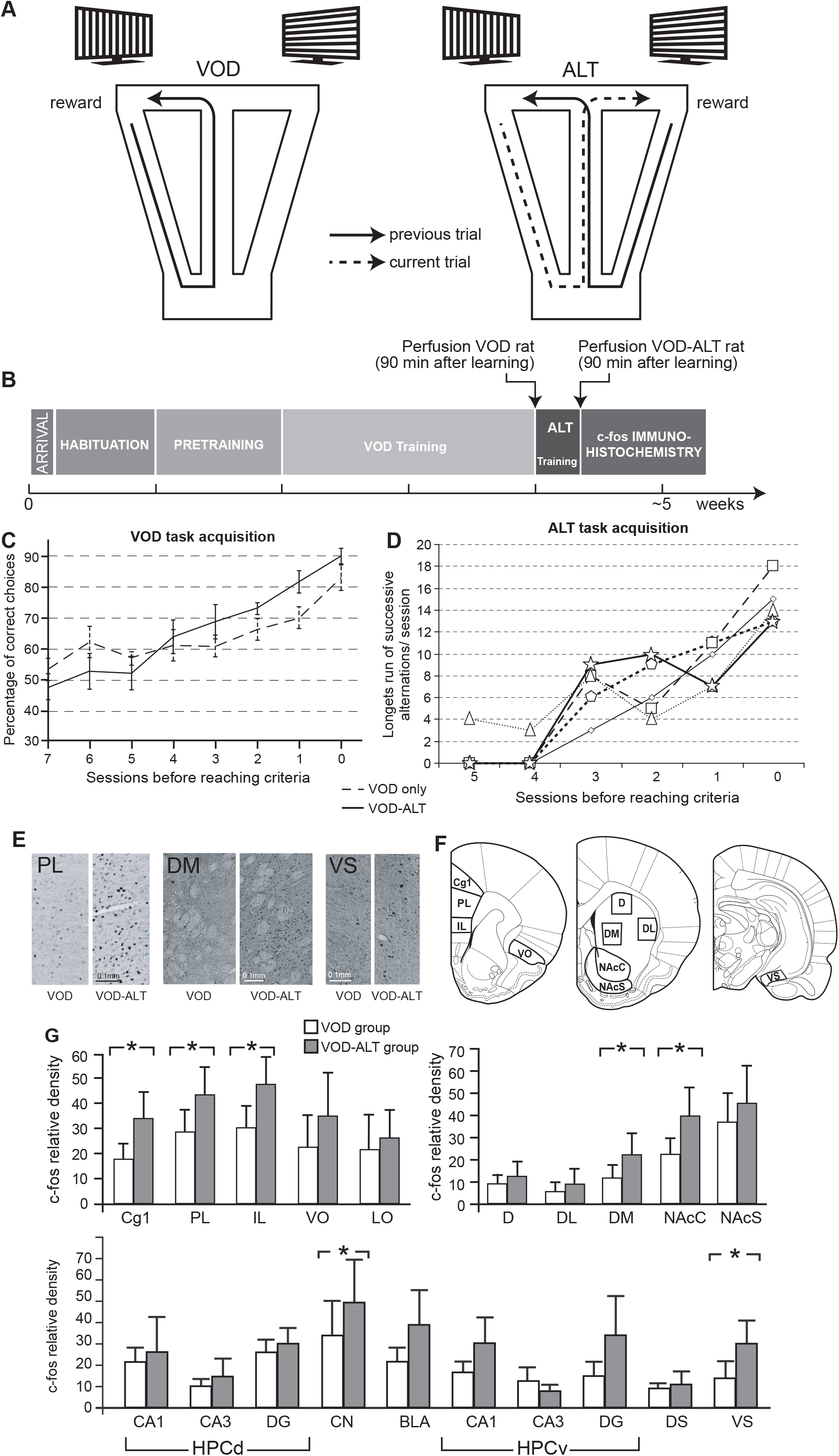
(A) As the rat crossed the middle of the central arm of the T-maze with return arms, cues appeared on the two screens. Correct choices were rewarded for going to the arm with vertical stripes in the VOD task and for going to the opposite arm from the previous trial in the ALT task. (B) Time course of experimental protocol for each cohort of rats. Learning curves for the respective rats for acquisition of (C) the VOD task and (D) the ALT task. (E) Representative histological sections showing immunoreactive neurons. (F) Sites sampled for neuron counts (adapted from Paxinos and Watson, 1998). (G) Relative density of c-fos immunoreactivity. Stars indicate p<0.05 significance in a one-tailed paired t-test for the five cohorts of rats. Abbreviations: Cg1 – cingulate area 1; PL – prelimbic area; IL – infralimbic area; VO – ventral orbital cortex; LO – lateral orbital cortex; D – dorsal striatum; DL – dorsolateral striatum; DM – dorsomedial striatum; NAcC and NAcS – nucleus accumbens core and shell; HPCd and v -dorsal and ventral hippocampus; CN and BLA – central and basolateral amygdalar nuclei; DS and VS – dorsal and ventral subiculum.

The rats were first habituated to the T-maze over three to five days by allowing them to forage for Kellogg’s Choco-Pops^®^ (which they had previously sampled in their home cages) that were distributed randomly along the surface. Habituation was terminated after the rats moved about easily and consumed food on all parts of the maze without freezing or defecating. Rats were then pretrained over three to nine days to run in the correct direction in the maze, starting from the base of the central arm, then turning left or right at the ‘T’, and proceeding to the reward site. At first, guillotine doors (controlled remotely by a manual pulley system) prevented backtracking after entries into the selected reward arm. During pretraining, a drop of saccharinated water was delivered for either choice. Then the rat had to continue back along the return arm located on the same side to start a new trial, and a movable barrier prevented direction reversal. If the rat turned in only one of the directions at the choice point, a barrier was put into place for several trials to force visits to the other side. Pretraining ended after the rats became proficient at performing moving in the correct direction without guidance.

The experiment was carried out in five replications, each with a cohort of two rats, trained in successive sessions and their brain tissues were processed together (Figure 1B). This permitted reduction of variability and hence a reduced sample size since all comparisons involved only pairs of rats exposed to identical housing and training conditions, and tissues were processed under identical conditions with identical reagents. They were first trained in a visual orientation discrimination (VOD) task. One rat of the cohort was then anesthetized and perfused while the other was trained in an alternation task (ALT) in the same T-maze. Rats were selected randomly for participation in the two respective groups. In the VOD task, the screen with the vertically oriented black stripes was behind the arm associated with reward on that trial, while in the ALT task rats were rewarded for choosing the reward arm opposite to the one selected on the previous (rewarded or unrewarded) trial, ignoring the visual cues. For unrewarded choices, the rats had to continue down the return arm to initiate a new trial. The first trial of ALT sessions was not rewarded. The onset of the ALT task was signaled by an intermittent tone (repetition of the Microsoft Windows standard system sound ‘Asterisk’) presented from a loudspeaker in front of the T-maze simultaneously with the visual cues. The tone continued until the rat completed a turn onto a reward arm in each trial. Otherwise, all stimuli remained the same as in the VOD task. The VOD-ALT group first learned the VOD task, and then was trained to criterion in the ALT task. Note that this experimental design emulates those employed in brain imaging studies. In order to model the neurophysiological recording protocol of Peyrache et al. (2009) and Benchenane et al. (2010) and to accelerate the training process, rats were permitted as many trials as they would run in a given session, and up to three sessions were run per day. Sessions were stopped when the animal stopped moving and all sessions lasted less than 30 min. Intervals between sessions were at least 3 hours.

To establish the learning criterion for the VOD task, we counted the number of runs of 4 or more successive correct trials, corresponding to a binomial probability of p≤(0.5)^4^, or p≤0.0625. The total number of trials in these runs had to exceed 50% over two consecutive days. Alternatively, a total of 80% correct trials (irrespective of run length) in a single session was also considered to satisfy the learning criterion (see Figure 1C). The rat was perfused 90 min later since this was reported to be the optimal delay for revealing activation-dependent c-fos accumulation (Sheng and Greenberg, 1990). The criterion of VOD performance for VOD-ALT group prior to ALT training was 80% correct (irrespective of run length) over 50 successive trials (which could be from the last two sessions, if the final one was too brief). The learning criterion for the ALT task was a run of 12 or more alternation trials in a given session, and again a 90 min delay preceded perfusion. These elevated criteria were necessary to assure reliable task performance.

Rats were administered a lethal dose of pentobarbital and perfused with saline followed by freshly prepared 4% paraformaldehyde in phosphate buffer (PB, pH 7.4). Brains were removed and post-fixed in the buffered 4% paraformaldehyde solution for 24 hours, then placed in PB containing 30% sucrose (pH 7.4). Frozen sections were cut coronally at 40 μm. Sections from brains of the two rats belonging to the same cohort (one VOD rat and one VOD-ALT rat) were processed simultaneously. Sections were first incubated in 0.3% H_2_O_2_ for 30 min. After three rinses, the sections were incubated overnight at room temperature in PB containing 0.2% primary c-fos antibody (sc-52, Santa Cruz Biotechnology, Dallas, TX, USA), 0.5% Triton X-100 and 0.1% bovine serum albumin (BSA; Sigma). After three rinses, the sections were incubated during 2 h in PB containing 0.5% secondary biotinylated antibody, 0.5% Triton X-100 and 0.1% BSA. After 4 rinses, the sections were incubated in avidin-biotin-peroxidase complex (ABC, Vector Laboratories, Burlingame, USA) solution for 2 h. Finally c-fos immunoreactivity was revealed with diaminobenzidine (DAB; 0.05%).

Regions of interest were carefully delineated and verified on adjacent sections treated with Nissl stain. C-fos immunostained neurons were counted automatically with Fiji image processing software (http://fiji.sc/Fiji) by two observers blind to treatment groups. Paired t-tests were used for statistical evaluation, with p<0.05, one-tailed, as the level of significance. One-tailed tests were used since the goal of the study was to determine the extent of the network with increased activity after the new acquisition relative to the initial acquisition.

All experiments were approved by the local animal experimentation ethics committee (Comité d’Ethique en Matière d’Expérimentation Animale-Paris Centre et Sud 59), and in accord with international (Directive 86/609/EEC; ESF-EMRC position paper 2010/63/EU; NIH guidelines) standards and legal regulations (Certificate no. 7186, Ministère de l’Agriculture et de la Pêche) regarding the use and care of laboratory animals.

## Results

### Behavioral performance

The VOD group reached performance in 17.8±4.5 (SEM) sessions, comprising 759±240 trials over 4 to 12 training days. The VOD-ALT group reached VOD performance criterion in 13.2±4.7 sessions comprising 634.4±210.4 trials. The number of trials and days to VOD criterion in the VOD and VOD-ALT groups was not significantly different (two-way paired Student t test, p=0.53 for trials and p=0.30 for days). Training for criterion performance in the ALT task required 4.2±0.36 sessions comprised of 311±20.5 trials (see Figure 1D).

During training in both tasks, the rats spontaneously used other strategies (such as go left, go right, ALT during the VOD task and VOD during the ALT task) despite the fact that this led to rewards on only 50% of the trials, on average. The principal tendency was for spontaneous alternation during the VOD training. To evaluate this tendency, only runs of six or more successive visits to the arm opposite the one visited on the previous trial were counted (binomial probability p≤0.0325) and these constituted 30±7% of the trials during training in the VOD group.

As expected, during ALT training, the rats all initially persisted in the VOD strategy. In all cases, they performed a minimum of 6 successive VOD trials in the penultimate or final session when the ALT criterion was reached. This indicates that rule shifting had indeed taken place since the most recent performance of the VOD strategy was in the recent past when they acquired the ALT strategy.

### c-fos labeling

The VOD-ALT group had significantly higher levels of c-fos immunoreactivity relative to the VOD group in the cingulate (Cg1), prelimbic (PL) and infralimbic (IL) cortices (respectively t(4)=2.58, t(4)=2.40, t(4)=2.48, pairwise Student t test, one-tailed, p<0.05, Figure 1E, *left*). No significant increase was detected in ventral orbital (VO) or lateral orbital (LO) areas. In the striatum, significantly higher levels of c-fos immunoreactivity were observed in the dorsomedial striatum (DM) and the core of the nucleus accumbens (NAcC) (respectively t(4)=2.67, t(4)=2.19; pairwise Student t test, p<0.05, one-tailed, Figure 1F). However, no such increases appeared in the dorsal (D) or dorsolateral (DL) striatum or in the accumbens shell (NAcS).

Burnham et al (2010) observed that c-fos immunoreactivity levels are closely related to the number of trials performed on the final training day. Here, the number of trials in the VOD and ALT groups on the final testing day did not show a significant difference (two-tailed paired t test p=0.22), and thus this is unlikely to be a possible confound.

In the hippocampal formation (not shown), there was a remarkably low density of c-fos immunoreactive neurons with no significant differences between the two groups in dorsal or ventral CA1, CA3 or dentate gyrus. Despite the small number of marked neurons, ventral subiculum c-fos immunoreactivity was significantly higher in the VOD-ALT group (t(4)=3.39; pairwise Student t test, one-tailed, p<0.05, not shown) while there was no difference between the groups in the dorsal subiculum. A significantly higher level of c-fos for the VOD-ALT group also appeared in the central nucleus of the amygdala (t(4)=2.30; pairwise Student t test, p<0.05, one-tailed, not shown) while the difference did not reach significance in the basolateral nucleus of the amygdala.

Note that although not statistically significant, the mean values of immunoreactive relative density were higher in the VOD-ALT group than the VOD group in all but one of these hippocampal system structures (the exception was ventral CA3, which also had extremely low densities). Brainstem neuromodulatory centers were also examined (substantia nigra pars compacta and reticulata, dorsal raphé nucleus, laterodorsal tegmental nucleus, ventral tegmental area, raphé nucleus, dorsal tegmental nucleus and locus cœruleus) – all had very low relative densities of marked neurons. In those with sufficient staining to permit analyses, there were no significant differences between the two groups (not shown).

## Discussion

The principal findings here are that the number of c-fos immunoreactive neurons increases in prefrontal cingulate, prelimbic and infralimbic areas, dorsomedial striatum and the core of the nucleus accumbens, as well as ventral subiculum and the central nucleus of the amygdala in animals learned a new rule compared to animals learning the previous rule. There are anatomical connections among these zones (Fudge *et al*., 2002; Jay and Witter, 1991; Mailly *et al*., 2013), and thus they are likely to compose functional networks. The absence of significant differences in the hippocampus is consistent with previous neurophysiological recordings of the hippocampus and nucleus accumbens of rats in a plus-maze as they alternated between spatial orientation strategies employing distal cue configurations vs. visible beacons. In that study we observed that hippocampal neurons do not change their firing fields between the two tasks (Trullier *et al*., 1999), but that nucleus accumbens neurons do change their firing patterns after task changes and platform rotations (Shibata *et al*., 2001). Thus in these studies the rule shifting was reflected by changes in accumbens neural activity but not hippocampal activity.

A popular experimental paradigm for testing task switching is to train rats in ‘place’ and ‘response’ rules in a plus-maze. The place task rewards visits to a specific arm while the response strategy requires the same body turn at the choice point. Rich and Shapiro (2007) inactivated PL/IL with muscimol injections and, in contrast with the present results, they observed that task switch acquisition was not impaired, but memory for the recently acquired switch was indeed impaired 24 h later. Instead, the rats persevered at the initially learned strategy. However, Ragozzino *et al*. (1999a) did observe switching impairments with reversible inactivation of PL/IL by local tetracaine infusion. Similar impairments in rule shifting were observed by Oualian and Gisquet-Verrier (2010) after PL and/or IL infusions with ibotenic acid. A possible explanation for this discrepancy advanced by Rich and Shapiro (2007) would be that alternate networks would support this acquisition. Our data would suggest that Cg1 could be part of such a network.

Burnham *et al*. (2010) examined c-fos immunoreactivity in rats acquiring an extradimensional shift between rules in a digging task with odor and texture cues. They found increased activity relative to cage controls in medial prefrontal cortex and orbitofrontal cortex. In a study of male F-344 rats shifting between response and place strategies in the plus maze, Grella *et al*. (2013) observed more activation of the immediate early gene *arc* in PL, IL and VO (but not MO and LO) than in cage control animals. While the tendency for increased c-fos activity in VO in VOD-ALT animals did not reach significance here, this does not preclude its participation in rule shifts. The use of cage-mate controls in the latter studies did not control for the increased sensory and motor activity of the maze training. The present results thus further demonstrate that after shifts between rules, there are also selective increases relative to animals having the same sensorimotor experience, and having reached criterion on a rewarded task (thus controlling for activation of reward circuitry).

A possible confound in interpreting the present data would be that the higher levels of c-fos activity in the VOD-ALT group might be related to the greater working memory requirement of the ALT task rather than to the rule shift. However, during VOD training, the VOD-ALT group showed a substantial tendency for spontaneous alternation on about 30% of training trials. Thus, the rats were already capable of using a strategy dependent upon working memory; the crucial difference is that they learned that this behavior led to reliable reward only when they reached the ALT criterion performance level. Perhaps the circuit reconfiguration associated with this learning leads to the c-fos activation. This would be consistent with our previous electrophysiological observations of changes in circuit properties in the PL upon new learning of a rewarded rule: hippocampal inputs are more effectively transmitted by inhibitory interneurons to principal neurons, and the latter change their phase of firing relative to hippocampal theta oscillations. Synchronously active cell assemblies are then formed, likely in relation to dopaminergic neuromodulatory inputs (Benchenane, *et al*., 2010). Thus, the reward expectation that comes with actual learning of the behavior-reward contingency would be expected to bring about changes in the network underlying the behavior, and this would be reflected in the c-fos activity.

The present results show, for the first time, striatal activation during acquisition of a new rule requiring selection among orienting cues and point to a prefrontal striatal-network underlying this behavioral and cognitive flexibility, consistent with lesion and pharmacological manipulation studies (Bissonette & Roesch, 2017). While this immediate early gene study identifies members of this network, electrophysiological studies will be required to determine with temporal precision the dynamic roles of these brain regions in switching attention from one stimulus modality to another and modifying behavior according to new reward contingencies.

## Acknowledgements

Thanks to France Maloumian for help preparing figures, and to Marie-Annick Thomas and Suzette Doutremer for training and advice for histology. Nicole Quenech’du and Jérémie Teillon for advice and training for image processing and Drs. Anne Cei and Michaël Zugaro for comments on the manuscript. Support came from French National Research Agency ANR-2010-BLAN-0217-01 Neurobot. A.B. was supported by the Erasmus Program of the European Community. Thanks to MEMOLIFE Laboratory of Excellence, Fondation Bettencourt Schueller.

## Conflict of Interest Statement

The research was conducted in the absence of any commercial or financial relationships that could be construed as a potential conflict of interest.

## Author Contributions

SW, HYG and VO developed the experimental design, SJS supervised the initial experiments, HYG, VO and AB performed the experiments, all authors analyzed the data, SW, HYG, VO and SJS wrote the manuscript, and all authors approved the final version.

## Notes

### Competing Interest Statement

The authors have declared no competing interest.

### Summary of Updates

Figure legend has been corrected.

